# The innovation of the symbiosome has enhanced the evolutionary stability of nitrogen fixation in legumes

**DOI:** 10.1101/2022.03.04.482842

**Authors:** Sergio M. de Faria, Jens J. Ringelberg, Eduardo Gross, Erik J.M. Koenen, Domingos Cardoso, George K.D. Ametsitsi, John Akomatey, Marta Maluk, Nisha Tak, Hukam S. Gehlot, Kathryn M. Wright, Neung Teaumroong, Pongpan Songwattana, Haroldo C. de Lima, Yves Prin, Charles Zartmann, Janet I. Sprent, Julie Ardley, Colin E. Hughes, Euan K. James

## Abstract

- Nitrogen-fixing symbiosis is globally important in ecosystem functioning and agriculture, yet the evolutionary history of nodulation remains the focus of considerable debate. Recent evidence suggesting a single origin of nodulation followed by massive parallel evolutionary losses raises questions about why a few lineages in the N_2_-fixing clade retained nodulation and diversified as stable nodulators while most did not. Within legumes, nodulation is restricted to the two most diverse subfamilies, Papilionoideae and Caesalpinioideae, which show stable retention of nodulation across their core clades.
- We characterize two nodule anatomy types across 128 species in 56 of the 150 genera of the legume subfamily Caesalpinioideae: 1) fixation thread nodules (FTs), where nitrogen-fixing bacteroids are retained within the apoplast in modified infection threads and 2) symbiosomes, where rhizobia are symplastically internalized in the host cell cytoplasm within membrane-bound symbiosomes.
- Using a robust phylogenomic tree based on 997 genes from 146 caesalpinioid genera, we show that losses of nodulation are more prevalent in lineages with FTs.
- We propose that evolution of the symbiosome allows for a more intimate and enduring symbiosis through greater compartmentalisation of their rhizobial microsymbionts, resulting in greater evolutionary stability of nodulation across this species-rich pantropical clade of legumes.

## Introduction

The N_2_-fixing clade of angiosperms includes all plants that form specialised organs known as nodules, within which they house intracellular diazotrophic bacteria (van Velzen *et al.*, 2018a). Within this clade, some species of Cucurbitales, Fagales and Rosales engage in nodulating symbiosis with the filamentous actinobacteria *Frankia*, while *Parasponia* (Rosales, Cannabaceae) and legumes (Fabales, Fabaceae) host phylogenetically diverse strains of *Alpha-* and *Betaproteobacteria* collectively known as rhizobia (Soltis *et al.*, 1995; Sprent *et al.*, 2017; Griesmann *et al.*, 2018; van Velzen *et al.*, 2018a). Strikingly, nodulation is mostly a rare trait across these four orders, having been reported in relatively few species except in Fabales, where the majority of the c. 20,000 species in the Fabaceae appear to be nodulated (Doyle, 2011). Across the legume family, nodulation is also very unevenly distributed, with most species in Papilionoideae and the Mimosoid clade (Caesalpinioideae *sensu* LPWG (2017)) being nodulated, whereas nodulation is less common in non-mimosoid Caesalpinioideae and absent in the other four smaller legume subfamilies (LPWG, 2017). The reasons for this uneven phylogenetic distribution of nodulation are unclear.

Despite the ecological and economic significance of N_2_-fixing root nodule symbiosis in ecosystem functioning and agriculture (Peoples *et al.*, 1995; Batterman *et al.*, 2013; Vitousek *et al.*, 2013; Epihov *et al.*, 2017), there is no consensus about the evolutionary origins of this important trait. Hypotheses have shifted from a scenario of multiple origins (Doyle, 2011; Werner *et al.*, 2014), potentially predisposed by a cryptic precursor that evolved in the ancestor of the N_2_-fixing clade (Soltis *et al.*, 1995; Werner *et al.*, 2014), to one of a single origin and massive parallel evolutionary losses (Griesmann *et al.*, 2018; van Velzen *et al.*, 2018a,b). This second hypothesis was generally dismissed because multiple independent origins provided a more parsimonious solution for the phylogenetic distribution of nodulating lineages, and because variation in nodule types and microsymbionts suggested that nodules are potentially non-homologous and arose multiple times. This clustered homoplasious occurrence of nodulation, confined to just one clade of angiosperms (Marazzi *et al.*, 2012), prompted the idea that a cryptic precursor evolved in the ancestor of the N_2_-fixing clade, which conferred a propensity for nodulation that was expressed in just a subset of lineages (Soltis *et al.*, 1995; Doyle, 2011, 2016; Werner *et al.*, 2014). However, no evidence for such a precursor, genetic or otherwise, has been found (Doyle, 2016; Griesmann *et al.*, 2018; van Velzen *et al.*, 2018a). Furthermore, nodulation involves structural and biochemical innovations underpinned by many genes, multiple developmental and signalling pathways, and coordination between the host and the microsymbiont (Brewin, 2004; Oldroyd & Downie, 2008; Oldroyd, 2013; Sprent *et al.*, 2017; Ardley & Sprent, 2021; Ledermann *et al.*, 2021), such that evolutionary gains of nodulation are likely to be more difficult than losses (van Velzen *et al.*, 2018a; Edwards, 2019). Recently, the alternative hypothesis of a single evolutionary origin of nodulation followed by numerous parallel evolutionary losses has gained traction, notably from comparative genomic studies documenting pseudogenization or loss of key nodulation genes in non-nodulating species, indicative of secondary losses of nodulation (Griesmann *et al.*, 2018; van Velzen *et al.*, 2018a; Zhao *et al.*, 2021). Re-examination of the structural and developmental homologies and commonalities in symbiotic gene function across nodulating lineages spanning the N_2_-fixing clade suggested that these also provide more compelling evidence for the single gain and multiple losses hypothesis (van Velzen *et al.*, 2018a).

This shift in thinking prompts questions about how the numerous secondary losses of nodulation are distributed across lineages and through time, why certain lineages retained nodulation to diversify as stable N_2_-fixers whereas many others lost this trait, and why, through time, N_2_-fixing symbiosis apparently became non-advantageous for the large majority of N_2_-fixing clade lineages.

One trait that has not been considered as a potential determinant of evolutionary stability of nodulation is the occurrence of two distinct anatomical arrangements of N_2_-fixing bacteria within the nodule. In the majority of papilionoid legumes, such as pea and *Medicago*, an infection thread (IT), formed from invagination of a root hair cell wall, conveys rhizobia from the point of infection to the nodule primordium. Rhizobia within the IT are budded off once they reach the nodule cell and are retained within it only by the host plasmalemma-derived symbiosome (or peribacteroid) membrane, where they differentiate into their N_2_-fixing bacteroid forms (Sprent, 2001, 2009; Brewin, 2004; Sprent *et al.*, 2017; Parniske, 2018; Ardley & Sprent, 2021; Tsyganova *et al.*, 2021). In contrast, in all actinorhizal symbioses and in a subset of nodulating legumes the N_2_-fixing bacteria are retained within modified, thin-walled, infection threads called fixation threads (FTs), remaining enclosed within the plant cell wall and the plasmalemma. Hereafter, we refer to these as FT--type nodules, and those in which the bacteroids are enclosed in symbiosomes as SYM-type nodules.

FT-type nodules were first described in actinorhizal plants (enclosing their *Frankia* microsymbionts), and in *Parasponia*, the only non-legume known to form nodules with rhizobia (Trinick, 1980; Lancelle & Torrey, 1985; Smith *et al.*, 1986). They were later observed in legumes, mostly in woody Caesalpinioideae (*sensu* LPWG (2017)) where they appeared to be relatively common (de Faria *et al.*, 1986, 1987; Naisbitt *et al.*, 1992; Sprent, 2001; Fonseca *et al.*, 2012). FT-type nodules have also been reported in a few legumes belonging to subfamily Papilionoideae, which is sister to Caesalpinioideae (Koenen *et al.*, 2020b; Zhao *et al.*, 2021), including tree genera such as *Andira*, *Dahlstedtia* and *Hymenolobium*, and members of tribe Brongniartieae (de Faria *et al.*, 1986, 1987; Sprent, 2001, 2009; Sprent *et al.*, 2013, 2017). Ultrastructural and histochemical analyses of FT-type nodules in *Parasponia* with rhizobia (Smith *et al.*, 1986), actinorhizal nodules with *Frankia* (Pawlowski & Demchenko, 2012), and in some legumes (de Faria *et al.*, 1986, 1987; Naisbitt *et al.*, 1992), revealed that FTs are superficially similar to the cell wall-bound “invasive” IT *e.g.* in harbouring some pectin (Fonseca *et al.,* 2012). The IT is an extension of the host cell wall and comprises mainly cellulose and pectin; its role appears to be largely protective, preventing the bacteria from invading the plant in a disorganised or pathogenic manner (Brewin, 2004; Tsyganova *et al.*, 2021). However, the composition and role of the FT remain uncertain. Moreover, the precise nature of the FT in relation to the symbiosome membrane, which in SYM-type nodules is essential for the exchange of nutrients between the host cytoplasm and the bacteroid (White *et al.*, 2007), remains unknown.

Within legumes, nodulation is restricted to the two largest subfamilies, Caesalpinioideae (including the Mimosoid clade, formerly Mimosoideae) and Papilionoideae (Sprent *et al.*, 2017; Ardley & Sprent, 2021). Here we investigate the occurrence of FT- and SYM-type nodulation across Caesalpinioideae, the second largest subfamily of legumes, with 151 genera and c. 4600 species distributed pantropically across all lowland tropical biomes, with minor incursions into temperate regions. We provide an updated census of nodulation occurrence, including three new records in genera of previously unknown nodulation status, plus extensive new data about FT- and SYM-type nodules across genera. We investigate whether these two nodule types represent different degrees of ‘compartmentalisation’ *sensu* Chomicki *et al.* (2020), by examining the anatomy and structure of nodules in *Chidlowia*, *Pentaclethra* and *Erythrophleum*, genera that span one of two evolutionary transitions from FT-type to SYM-type nodules that we hypothesize to have occurred within Caesalpinioideae along the branch subtending the Mimosoid clade. *Erythrophleum*, which is sister to the Mimosoid clade, has FT-type nodules, while *Pentaclethra* and *Chidlowia*, as well as all other studied taxa in the Mimosoid clade (Manzanilla & Bruneau, 2012; Koenen *et al.*, 2020a), have SYM-type nodules.

We test the hypothesis that the transition in nodule anatomy from FT- to SYM-type nodules constituted an evolutionary innovation that led to more stable retention of nodulation, whereas lineages in which FT-type nodules occur are more prone to evolutionary losses of nodulation. For this, we explore the number and phylogenetic distribution of evolutionary losses of nodulation using a robust phylogenomic backbone that includes 97% of Caesalpinioideae genera. We also examined the composition of the FT wall in more detail than has hitherto ben achieved using immunohistochemical methods that have been used for ITs (Brewin, 2004; Tsyganova *et al.*, 2021) in order to better elucidate its possible role in symbiosis.

## Materials and Methods

### Nodulation and nodule anatomy

Basic nodulation data (nodulated or non-nodulated) were obtained from Sprent (2001, 2009), from papers or reports published since 2009, and from previously unpublished records in the databases of the authors, including new reports of nodulation status (Tables S1 and S2). Where no data are available the nodulation status of a genus is listed as Uncertain (Un) (Table S1).

Anatomical types (FT or SYM) were determined from published data for the specific taxa that were used to construct the phylogeny, or related species in the same genus (based on substantial data that indicate that nodulation is almost always a generic trait (Sprent *et al.*, 2017)), alongside extensive data newly obtained here (Tables S1 & S2). Samples were prepared for light and electron microscopy according to de Faria *et al.* (1986, 1987) and Fonseca *et al.* (2012), unless otherwise stated. Additional samples of nodules from *Chidlowia sanguinea*, *Entada polystachya*, *Erythrophleum* spp., *Moldenhawera* spp., and *Pentaclethra macroloba* were prepared specifically to examine in detail the presence of FTs or symbiosomes (Table S2). For these samples, slices from four or more nodules per species were fixed in 2.5% glutaraldehyde and processed in two ways: (1) for light microscopy and immunogold transmission electron microscopy (TEM) with the monoclonal antibody JIM5, which recognises unesterified pectin (VandenBosch *et al.*, 1989; Tsyganova *et al.*, 2021), according to Fonseca *et al.* (2012), and (2) for identifying the symbiosome membrane by TEM using additional post-fixation in osmium tetroxide followed by embedding in epoxy resin according to Rubio *et al.* (2009). Ultramicrotomy, staining of sections for light microscopy and for TEM, and immunogold labelling with JIM5 for TEM were as described in Fonseca *et al.* (2012).

For immunohistochemical analysis of the FT wall confocal laser scanning microscopy (CLSM) was performed on slides containing semi-thin sections (1 µm thickness) of *Erythrophleum* and *Pentaclethra* nodules fixed and embedded as per method (1) above. The sections were incubated for 2 h in 1:10 dilutions of monoclonal antibodies raised against various plant cell wall components (all obtained from Plant Probes, Centre for Plant Sciences, University of Leeds, UK): Lm2, which labels ß-linked-GlcA in arabinogalactose protein (AGP) glycan; Lm5, which labels the pectic polysaccharide rhamnogalacturonan; and Lm15, which labels the XXXG motif of the non-pectic, non-cellullosic polysaccharide xyloglucan. Then, after washing twice in distilled water (dH_2_O) the sections were incubated for 1 h in 1:500 dilution of goat anti-rat Alexa 488 secondary antibody (ThermoFisher, Loughborough, UK) followed by several rinses with dH_2_O. After mounting in coverslips and Fluoromount (ThermoFisher, Loughborough, UK), the sections were examined using a Zeiss LSM 710 confocal laser scanning microscope (Carl Zeiss Microscopy Limited, Cambourne, UK), fitted with a W Plan-Apochromat 40x lens, using spectral imaging with excitation at 488 nm and emissions between 494 nm and 727 nm. The images were colour-coded according to wavelength and enhanced using the Min/Max function in Zen 2010 software. Ultrathin sections (80 nm) of the same samples were then immunogold labelled for TEM using the same monoclonal antibodies as those for CLSM (Lm2, Lm5, Lm15) according to Fonseca *et al.* (2012).

### Phylogeny and ancestral trait estimation

We used a recently constructed time-calibrated phylogeny of Caesalpinioideae that included 146 of the 151 genera, which was based on targeted enrichment of 997 nuclear genes using the Mimobaits gene set (Koenen *et al.*, 2020a) to generate a large phylogenomic Hybseq dataset (Ringelberg et al., unpublished). Using this much larger gene set allowed us to overcome lack of resolution prevalent across the backbone of the non-mimosoid grade in previous phylogenies that were based on traditional Sanger DNA sequence datasets (Bruneau *et al.*, 2008; Manzanilla & Bruneau, 2012; LPWG, 2017) to generate a robust and densely sampled phylogenetic hypothesis. The five unsampled genera are *Hultholia*, a member of the Caesalpinia clade (Gagnon *et al.*, 2016), a group of 27 genera that are all either non-nodulating or of unknown nodulation status; *Stenodrepanum*, which is sister to and doubtfully distinct from *Hoffmannseggia*, and is also placed in the Caesalpinia clade (Gagnon *et al.*, 2016); *Pterogyne*, which is also non-nodulating and likely forms a phylogenetically isolated monogeneric lineage in the non-mimosoid grade of Caesalpinioideae that is potentially sister to a large clade comprising all Caesalpinioideae except the Umtiza and Ceratonia clades (Zhao *et al.*, 2021); *Microlobius*, which is nodulating with SYM-type nodules and likely nested within the genus *Stryphnodendron* (Simon *et al.*, 2016; Ribeiro *et al.*, 2018), and finally the non-mimosoid *Vouacapoua*, which is non-nodulating, but is likely placed in the Cassia clade which contains both nodulating and non-nodulating lineages (Bruneau *et al.*, 2008).

The original 420-taxon Caesalpinioideae phylogeny was time-calibrated in BEAST (Drummond & Rambaut, 2007), using a species tree topology estimated by ASTRAL (Zhang *et al.*, 2018) based on gene trees of 821 single- or low-copy genes (Ringelberg et al., unpublished). We trimmed this chronogram until each genus was represented by just a single taxon, with two exceptions. First, the genus *Chamaecrista*, for which we retained four species due to known variation in nodule type within that genus (Naisbitt *et al.*, 1992; Santos *et al.*, 2017). Second, for non-monophyletic genera (Ringelberg et al., unpublished) we retained representative taxa for each para-/polyphyletic lineage. Nodulation data (Table S1) were matched to the tips of this tree in as conservative a way as possible. For example, in the case of *Prosopis*, which is polyphyletic and is thus represented by four taxa in the tree, these were scored as follows: *P. juliflora*, *P. cineraria*, and *P. africana*, which represent three independent lineages, are scored as SYM, as the nodule type of these taxa is known. In contrast, the taxon representing the fourth lineage, *P. ferox*, is scored as nodulating but with an unknown nodule type, because *P. ferox* and other taxa in this clade are nodulating, but their nodule types are unknown. The extensive generic non-monophyly visible in Caesalpinioideae, especially within the mimosoid clade (Fig. 1) is the focus of current taxonomic work reported elsewhere (Ringelberg et al., unpublished).

**Fig. 1.**
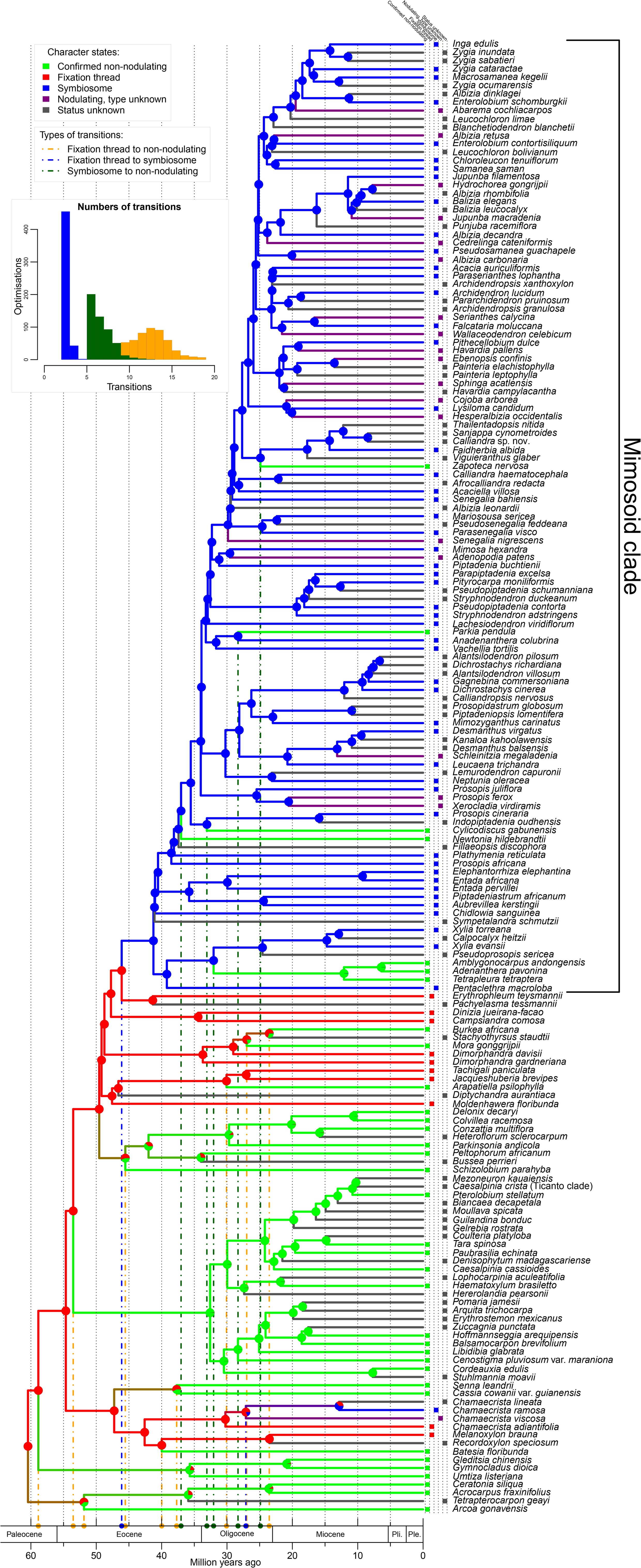
Evolutionary trajectories of nodulation and nodule type across a time-calibrated phylogeny of the Caesalpinioideae legumes. Pie charts on nodes show the proportions of the most likely reconstructed character states: non-nodulating, fixation thread (FT-type nodules), symbiosome (SYM-type nodules), nodulating but of unknown type, and nodulation status unknown, summarised over 500 simulations. Branch colours denote the nodulation status of the node or tip it subtends and the coloured boxes in front of each taxon name show the character state for that species. Note that in three clades (the *Senna* + *Cassia* clade, the *Arcoa* + *Acrocarpus* clade and the *Peltophorum* clade) ‘double’ losses of FT-types nodules are inferred to have occurred simultaneously in both descendant lineages of that node. For example, the crown node of the *Senna* + *Cassia* clade is inferred to be nodulating with FT-type nodules even though *Senna* and *Cassia* are non-nodulating. The dashed orange, blue, and dark green vertical lines show the phylogenetic locations and maximum ages of the various character state transitions on the tree. Using the same colours, the histogram shows the frequencies of the number of transitions from FT to SYM (blue), from SYM to non-nodulation (green) and from FT to non-nodulation (orange) across 500 independent character estimations. Note that the three other character state transitions, from non-nodulating to FT or SYM-type nodules, and from SYM to FT, were not allowed under our model, and were therefore fixed at zero. Pli = Pliocene; Ple = Pleistocene.

Nodulation status was estimated across this phylogeny using a model with three character states: non-nodulating, fixation thread (FT), and symbiosome (SYM). We explicitly followed the single gain, multiple losses evolutionary model for the origins of nodulation, i.e., nodulation evolved only once in angiosperms and was subsequently lost many times. To do this, we constrained the model so that the root state of the tree was set to FT and allowed only three types of transitions: from FT to non-nodulating, from FT to SYM, and from SYM to non-nodulating. Any transitions from a non-nodulating to a nodulating state were thus not allowed. We used stochastic character mapping implemented in the function make.simmap in the PHYTOOLS R package (Revell, 2012) to simulate 500 independent evolutionary trajectories of nodulation across the time-calibrated phylogeny. Results were summarised onto this tree across all simulations, with transitions inferred along branches connecting nodes that have different character states in the majority of the 500 simulations. Taxa for which the nodulation status is unknown were assigned equal probabilities for all three character states.

## Results

### Nodulation and nodule anatomy

Data on nodule anatomy for 128 species of 56 genera of Caesalpinioideae, including more than 80 records newly reported here (Table S1, S2), show that no species belonging to the Mimosoid clade have FT-type nodules, i.e. all known nodulating mimosoids have SYM-type nodules (Fig. S1). In contrast, all nodulating species from the grade subtending the Mimosoid clade have FT-type nodules (Fig. S2), except for a subset of species of *Chamaecrista*. In that sense *Chamaecrista* is exceptional among the taxa of the non-mimosoid grade, as it harbours species with either FT or SYM-type nodules (Naisbitt *et al.*, 1992).

Mature nodules of *Erythrophleum ivorense* and *E. suaveolens,* placed in the sister group of the Mimosoid clade (Fig. 1), are indeterminate with a meristem at the tip and a large invasion zone (IZ) (Fig. 2a), containing cells being invaded by rhizobia (Fig. 2b), and an N2-fixing zone occupying most of the nodule volume; this contains a mix of infected and uninfected cells (Fig. 2a). Bacteria can be seen to invade the IZ cells via ITs emerging from between the cells (Fig. 2c); these ITs are intensely immunogold labelled with the monoclonal antibody JIM5 indicating that they contain unesterified pectin, specifically partially methylated homogalacturonan (VandenBosch *et al.*, 1989; Tsyganova *et al.*, 2021), and are similar in that respect to the host cell wall (Fig. 2c, d). Infected cells in the N2-fixing zone contain numerous FTs (Fig. 2e), bound by cell walls that are thinner than those of the ITs, and either have sparser labelling with JIM5 (Fig. 2d, f) or have no apparent labelling (Fig. 2f). Higher definition TEM with osmicated samples reveals that the FTs are surrounded by a cell membrane (Fig. 2g, h), that appears to be derived from the host endoplasmic reticulum (ER). This shows that the FT comprises a multi-layered compartment consisting of a cell membrane, the FT cell wall, the lumen of the FT, and the bacteroid (Fig. 2g). Infected cells of nodules on the neotropical legumes *Moldenhawera floribunda* and *M. blanchetiana* var. *multijuga*, also placed in the non-mimosoid grade of caesalpinioids (Fig. 1), are packed with bacteroids enclosed within FTs (Fig. S2a, b). As with *Erythrophleum*, the ITs were intensely labelled with JIM5 (Fig. S2c), but the FTs considerably less so (Fig. S2d); the FTs in *Moldenhawera* were also associated with membranes arising from the ER (Fig. S2e, f). Another three non-mimosoid neotropical caesalpinioid genera (*Jacqueshuberia purpurea*, Fig. S2g, h; *Tachigali rugosa*, Fig. S2i, j; and *Campsiandra comosa*, Fig. S2k, l), also have their bacteroids enclosed in FTs labelled to various degrees with JIM5. Together with published reports on *Chamaecrista* (de Faria *et al.*, 1987; Naisbitt *et al.*, 1992) and *Dimorphandra* (Fonseca *et al.*, 2012), these observations of five additional genera demonstrate the ubiquity of FT-type nodules across nodulating lineages in the non-mimosoid grade of Caesalpinioideae.

**Fig. 2.**
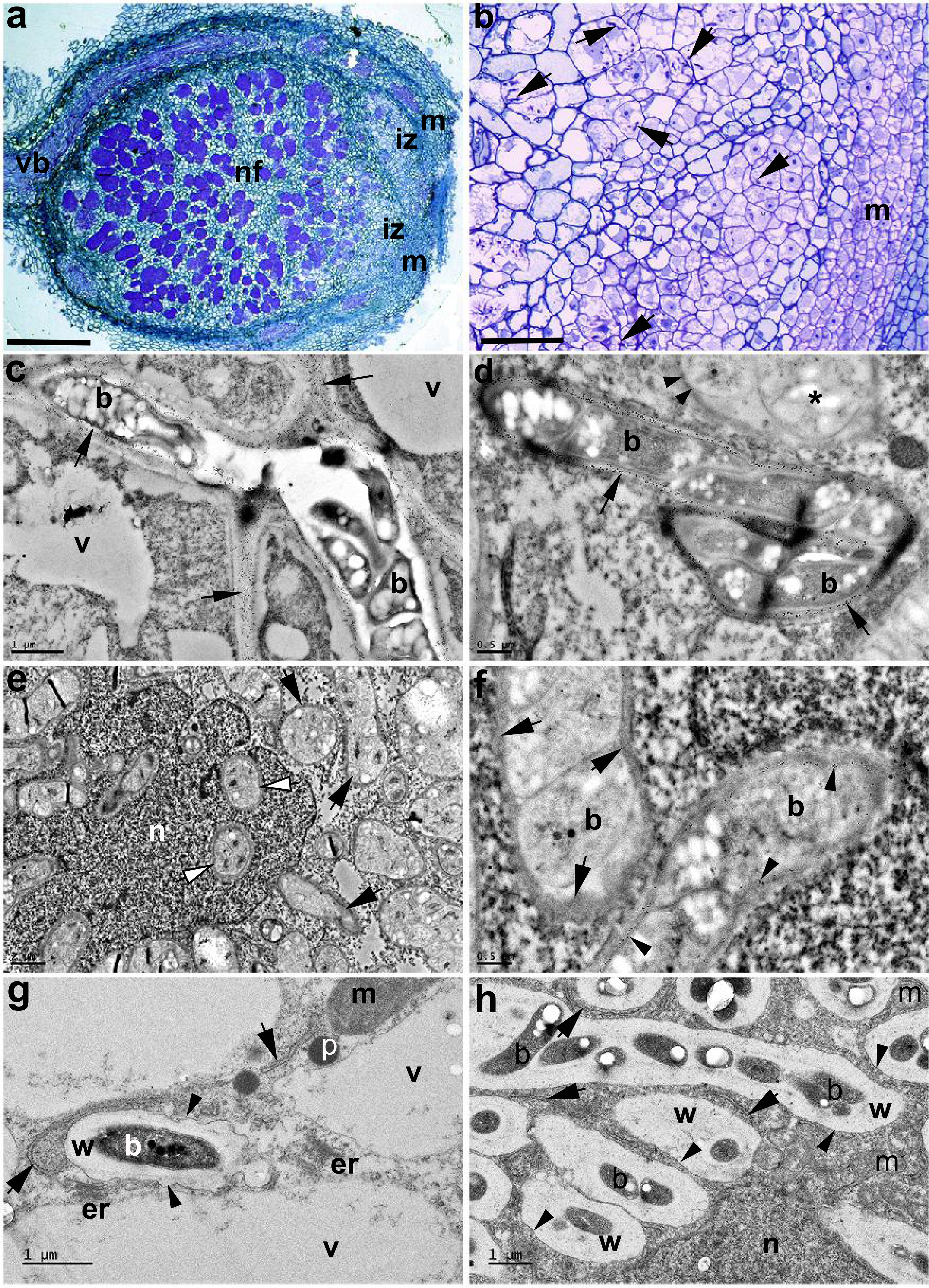
Non-mimosoid grade caesalpinioid nodules in the genus *Erythrophleum* contain bacteroids enclosed within fixation threads (FTs). Light (a, b) and transmission electron microscope (c-h) images of sections of nodules from *E. ivorense* (a-f) and *E. suaveolens* (g, h). **a,** whole nodule longitudinal profile illustrating the zonation typical of an indeterminate nodule (m = meristem, iz = invasion zone, nf = nitrogen fixing zone). Bar = 250 µm. **b,** higher magnification view of the iz in which newly-divided host cells derived from the meristem (m) are being invaded by numerous infection threads (arrows). Bar = 10 µm. **c,** large infection thread containing bacteria (b) invading cells in the iz; the walls of the infection thread are densely immunogold labelled with 10 nm gold particles linked to JIM5 (arrows), a monoclonal antibody which recognises non-esterified pectin. v = vacuole. Bar = 1 µm. **d,** infection thread in the iz-nf boundary with its cell walls labelled with JIM5 (large arrows) adjacent to an FT (*) with a thinner cell wall that is very sparsely labelled with JIM5 (arrowheads). Bar = 500 nm. **e,** cell in the nf zone packed with FTs (arrows), including within the nucleus (n). Bar = 2 µm. **f,** detail of FTs in the nf zone containing N-fixing bacteroids (b); the FT walls range from being sparsely labelled with JIM5 (arrowheads) to exhibiting little or no obvious labelling (single gold particles are indicated by arrows). Bar = 500 nm. **g,** high resolution image of a bacteroid (b) forming within a strand of cytoplasm between vacuoles (v) in an iz cell; the bacteroid is surrounded by a cell wall (w) that is being enveloped in a membrane (arrowheads), stretches of which (arrows) appear to be derived from nearby endoplasmic reticulum/Golgi bodies (er). The intense metabolic activity of this process is suggested by the nearby mitochondria (m) and peroxisomes (p). Bar = 1 µm. **h,** bacteroids in newly-formed FTs packed into a new N-fixing cell in the early nf zone adjacent to the iz; the bacteroids are surrounded by the FT wall (w), which is itself surmounted by a symbiosome membrane (arrowheads). Note the membranes within the cytoplasm that are associated with the FTs (arrows). n = nucleus, m = mitochondrion. Bar = 1 µm.

In contrast, *Pentaclethra macroloba* nodules are indeterminate (Fig. 3a) with infected cells in the N_2_-fixing zone surrounded by uninfected cells (Fig. 3b) containing bacteroids that are not surrounded by a cell wall (Fig. 3c) but are clearly enclosed within symbiosomes (Fig. 3d). In the same clade, *Xylia xylocarpa* also has SYM-type nodules (Fig. S1g, h). *Chidlowia sanguinea* nodules are similar to those of *P. macroloba*, except that the IZ is more prominent (Fig. 3e); the bacteroids are also enclosed in symbiosomes (Fig. 3f). *Chidlowia sanguinea* is sister to a large clade containing the bulk of mimosoid species, wherein SYM-type nodules are consistently present, as illustrated by *Entada polystachya* (Fig. S1a, b), *Enterolobium cyclocarpum* (Fig. S1c, d) and *Lachesiodendron viridiflorum* (Fig. S1e, f), showing that symbiosomes are universally found in nodulating lineages across the entire Mimosoid clade (Fig. 1, Table S1, S2).

**Fig. 3.**
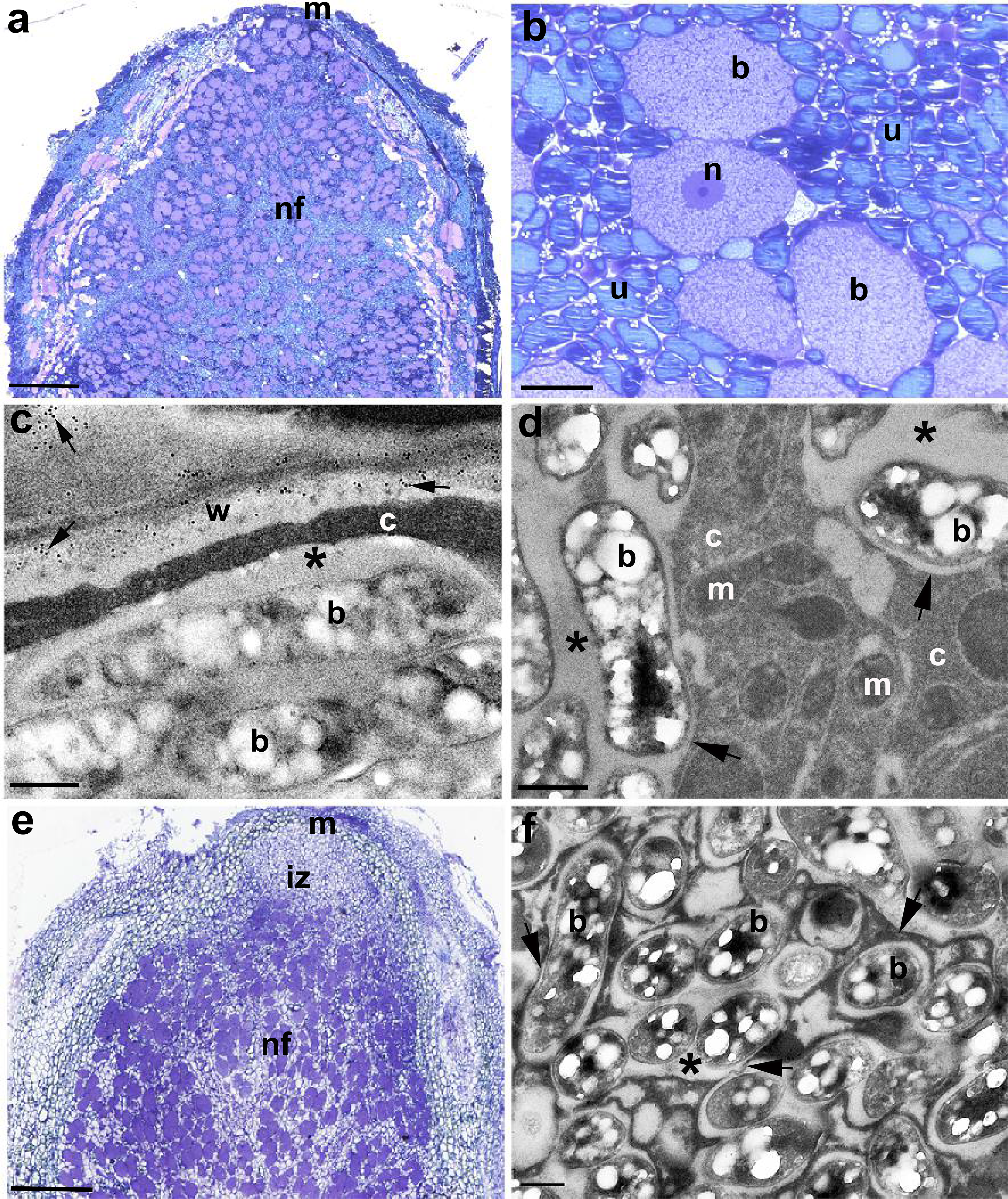
Nodules of caesalpinioids from the Mimosoid clade contain bacteroids enclosed within symbiosomes (SYMs). Light (a, b, e) and transmission electron microscope (c, d, f) images of sections of nodules from *Pentaclethra macroloba* (a – d) and *Chidlowia sanguinea* (e, f). **a,** whole *P. macroloba* nodule longitudinal profile illustrating the zonation typical of an indeterminate nodule (m = meristem, nf = nitrogen fixing zone). Bar = 500 µm. **b,** higher magnification view of the nf zone showing large bacteroid-containing cells (b) surrounded by smaller and more numerous uninfected cells (u). n = nucleus. Bar = 25 µm. **c,** bacteroids (b) within a symbiosome adjacent to the host cell wall (w) which is immunogold labelled with 10 nm gold particles linked to JIM5 (arrows). The symbiosome peribacteroid space is marked with *; note that there is no cell wall separating the symbiosome from the host cytoplasm (c). Bar = 500 nm. **d,** high resolution image of bacteroids (b) housed in symbiosomes; the symbiosome membrane separating it from the cytoplasm (c) are marked with arrows, and the peribacteroid space by *. m = mitochondrion. Bar = 500 nm. **e,** whole *C. sanguinea* nodule longitudinal profile illustrating the zonation typical of an indeterminate nodule (m = meristem, iz = invasion zone, nf = nitrogen fixing zone). Bar = 200 µm. **f,** high resolution image of bacteroids (b) housed in symbiosomes; the symbiosome membrane separating it from the cytoplasm (c) are marked with arrows, and the peribacteroid space by *. Bar = 500 nm.

The FT wall in *Erythrophleum* nodules was investigated further using monoclonal antibodies against arabinogalactose protein (AGP) glycan (Lm2), the pectic polysaccharide rhamnogalacturonan (Lm5), and the non-pectic, non-cellullosic polysaccharide xyloglucan (Lm15). These probes were capable of clearly delineating FTs in both confocal laser scanning microscopy (CLSM) (Fig. 4a – d) and TEM (Fig. 4e – h), indicating that the walls of FTs contain all three of these components. In contrast, the symbiosomes in *Pentaclethra* nodule sections treated identically were very difficult to discern using the CLSM (Fig. 4j – l), and although they could be observed under the TEM they had few or no gold particles associated with them indicating an absence of cell wall components (Fig. 4n – p). The exception was Lm2 (AGP glycan), which labelled *Pentaclethra* symbiosomes (Fig. 4i, m), and thus confirmed previous observations of AGP in the symbiosome membrane made with pea (*Pisum sativum* L.) nodules (Tsyganova *et al.,* 2021).

**Fig. 4.**
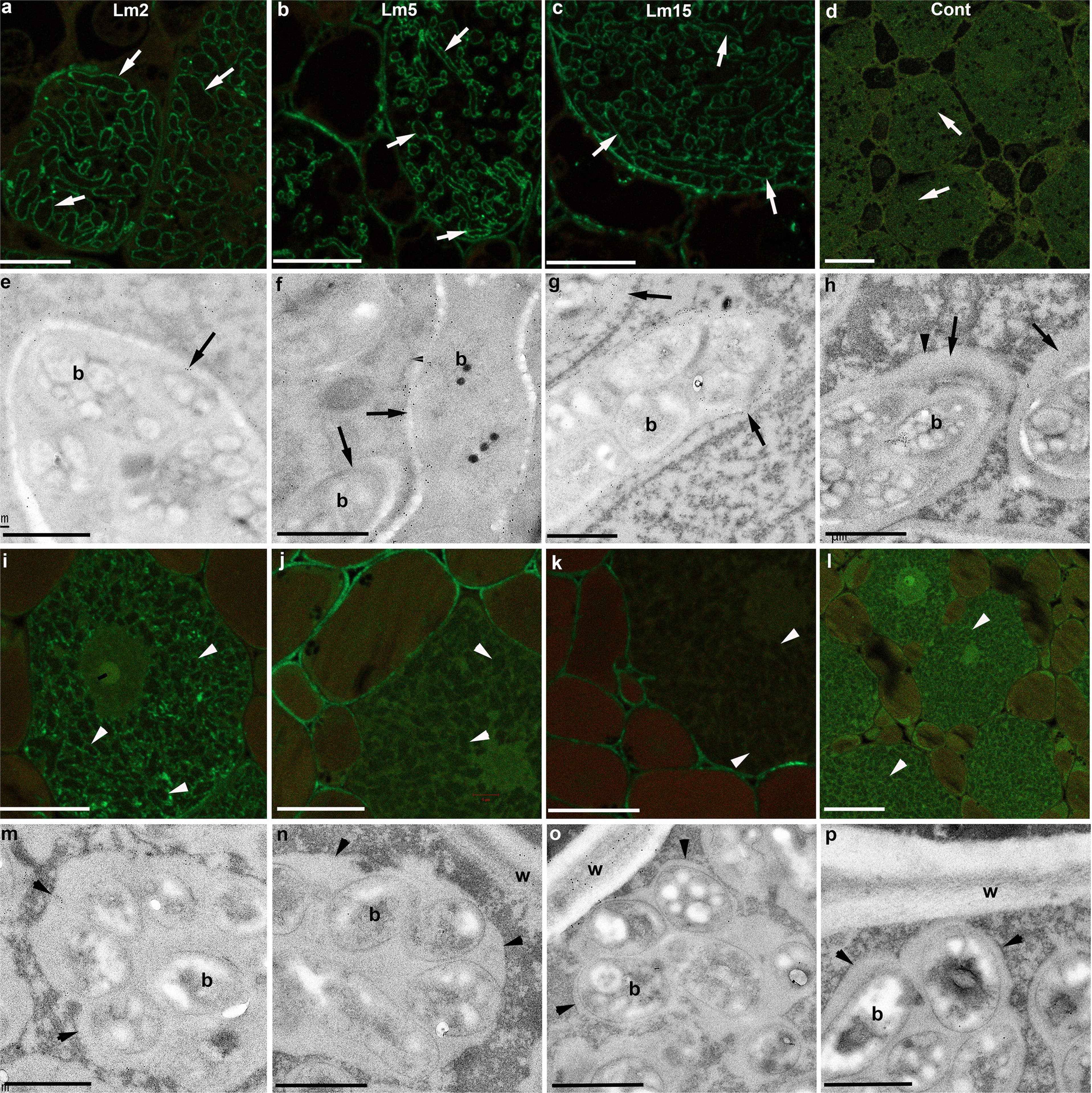
Fixation threads contain other cell wall components. Confocal laser scanning microscopy (CLSM) with anti-rat Alexa Fluor 488 (a – d, i – l) and immunogold TEM with anti-rat 10 nm gold (e – h, m – p) of *Erythrophleum* (a – h) and *Pentaclethra* (i – p) nodules incubated in monoclonal antibodies raised in rat against various plant cell wall components: Lm2 (a, e, i, m), which labels arabinogalactose protein (AGP) glycan; Lm5 (b, f, j, n) which labels the pectic polysaccharide rhamnogalacturonan; and Lm15 (c, g, k, o), which labels the XXXG motif of the non-pectic, non-cellullosic polysaccharide xyloglucan. Control sections incubated in buffer alone without a primary antibody are presented in (d, h, l, p). FTs are indicated by arrows in a – h, and symbiosomes by arrowheads in i – p. w = host cell wall separating plant cells, b = bacteroid. Bar = 5 µm (a – d, i – l), Bar = 1 µm (e – h, m – p).

### Phylogeny and evolution of nodule types

Ancestral estimation of nodulation and nodule types across the caesalpinioid phylogeny reveals two independent transitions from FT- to SYM-type nodules, first on the branch subtending the Mimosoid clade in the mid-Eocene, 45–40 Myr, and later within the genus *Chamaecrista* (Casaes et al. unpublished) in the early to mid-Miocene, 22–12 Myr (Fig. 1). Seventeen losses of nodulation are hypothesized across Caesalpinioideae, 12 in ancestrally FT-type, and five in ancestrally SYM-type lineages (Fig. 1, Table S3), suggesting that nodulation based on FT-type nodules is significantly more prone to loss than nodulation based on SYM-type nodules (Fig. 1). Exploratory analyses show that alternative scoring of character states for taxa with missing data, e.g. assigning equal weight to nodulation and non-nodulation states, does not significantly affect the outcome (Fig. S3).

Maximum ages of evolutionary losses of nodulation from FT-type nodule ancestry span the late Paleocene to the late Oligocene (59–24 Myr) and from SYM-type nodule ancestry from the late Eocene to the late Oligocene (37–25 Myr) (Fig. 1, Table S3).

## Discussion

Within legumes, nodulation is restricted to the two largest subfamilies, Caesalpinioideae (*sensu* LPWG (2017)) and Papilionoideae. The idea that nodulation was ‘stabilized’ in these two lineages was noted by Werner *et al.* (2014), who referred to core clades of Papilionoideae as ‘stable fixers’ i.e. a clade across which losses of nodulation were almost absent, and to the Mimosoid clade (Mimosoideae in Werner *et al.* (2014)) as having a ‘moderately stable fixing state’, where losses of nodulation were infrequent. Here we provide an explanation for this pattern: that the evolution of SYM-type nodules, with bacteroids free within a symbiosome, accounts for this greater stability. FT-type nodules characterize almost all nodulating non-mimosoid Caesalpinioideae genera, while the transition to SYM-type nodules on the stem lineage of the Mimosoid clade (Fig. 1) coincides with a shift to fewer losses of nodulation and greater stability of N_2_-fixation. Furthermore, the Mimosoid clade (c. 3500 species) is considerably more species-rich than the non-mimosoid grade (c. 1100 species), demonstrating a much higher number of losses of FT than SYM in a much lower number of species, i.e. a significantly lower rate of evolutionary losses per lineage per million years, further stressing the evolutionary stability of SYM compared to FT-type nodules (and/or higher diversification rates in SYM lineages).

In Papilionoideae, the sister group of Caesalpinioideae (LPWG, 2017; Koenen *et al.*, 2020b), nodulating and non-nodulating genera appear to be similarly intermingled phylogenetically across the initial divergences, while later stabilizing as nodulating across the core clades (Werner *et al.*, 2014; Doyle, 2016; Epihov *et al.*, 2017; van Velzen *et al.*, 2018a). It remains to be tested whether there is a similar association between more frequent losses of nodulation and FT-type nodules in Papilionoideae, where FT-type nodules are also found sporadically, but the vast majority of nodulating lineages and all the species-rich lineages – the ‘stable fixers’ (Werner *et al.*, 2014) – have SYM-type nodules. Lack of phylogenetic resolution among the initial divergences in Papilionoideae (Cardoso *et al.*, 2012, 2013; LPWG, 2017), mean that this test must await a more robust phylogeny.

The consistent occurrence of FT-type nodules across nodulating lineages in the non-mimosoid grade of Caesalpinioideae demonstrates that FT-type nodules are ancestral within Caesalpinioideae and persisted through early evolution of that subfamily (Fig. 1). The FT-type nodule also characterizes actinorhizal nodules (Pawlowski & Demchenko, 2012) and rhizobial nodules in *Parasponia* (Lancelle & Torrey, 1985) and is thus most likely ancestral across the N_2_-fixing clade (Shen & Bisseling, 2020) and legumes as a whole, where, as noted by Koenen *et al.* (2020b), rapid initial divergence of the six legume subfamilies implies additional losses of nodulation along the stem lineages or early in the crown group divergences of subfamilies Cercidoideae, Detarioideae, Dialioideae and Duparquetioideae, as no extant members of these subfamilies are known to nodulate. Although we have not looked at actinorhizal symbioses here, the very low proportion of nodulated species within Cucurbitales, Fagales and Rosales (Ardley & Sprent, 2021) further supports the hypothesis that lineages with FT-type nodules are more prone to evolutionary losses of nodulation.

It is well established that N_2_-fixation is energy-demanding, limited by photosynthesis, and confers fitness advantages only when nitrogen is limiting and when the benefits derived from greater availability of nitrogen, e.g. in fostering higher photosynthetic rates in N_2_-fixing plants, are greater than the costs of photosynthetic carbon (McKey, 1994; Hoffman *et al.*, 2014; Taylor & Menge, 2018; van Velzen *et al.*, 2018a). Additionally, there is experimental evidence showing that legumes increase N_2_-fixation at elevated CO_2_ levels and that nitrogenase activity declines rapidly above 35°C and below 25°C (Trinick, 1980), suggesting a greater advantage in being a N_2_-fixer under early Cenozoic CO_2_ levels and temperatures (Rogers *et al.*, 2009; Chen & Markham, 2021), and for those advantages to be preferentially retained in the tropics and subtropics, where FT-type nodulators are largely restricted. Falling atmospheric CO_2_ levels and temperatures through the Cenozoic could have triggered global evolutionary losses of nodulation across the N_2_-fixing clade (as suggested by van Velzen *et al.* (2018a)) and we show that maximum ages of losses of nodulation across Caesalpinioideae are widely scattered from the late Paleocene to the late Oligocene, 59 to 24 Myr (Fig. 1). However, it is important to consider that these are maximum ages rather than precise indicators of the timing of losses, and furthermore, the number of estimated losses per lineage is the minimum number of losses to explain the observed pattern at the tips, given the taxa sampled here.

It has long been recognized that evolutionary conflicts arise between hosts and symbionts over symbiont mixing, proliferation, and transmission (Frank, 1996) because the presence of multiple, genetically-heterogeneous symbiont strains within a host will cause symbionts to evolve traits that increase symbiont proliferation, competition, and conflict, but decrease the efficiency of the symbiosis (Frank, 1996). There is clear evidence that this happens in the rhizobia–legume symbiosis (Oono *et al.*, 2009; Sachs *et al.*, 2018). To resolve such conflicts, hosts have evolved ways to control symbiont proliferation (i.e., terminally differentiated bacteroids lose their ability to reproduce) (Mergaert *et al.*, 2006; De La Peña *et al.*, 2018; Ardley & Sprent, 2021), discriminate among symbionts (Yang *et al.*, 2017; De La Peña *et al.*, 2018; Ardley & Sprent, 2021), and penalise non-cooperating symbionts (Kiers *et al.*, 2003; Oono *et al.*, 2009; Ardley & Sprent, 2021). All these approaches are more effective when symbionts are compartmentalised within hosts, and all occur in the rhizobia–legume symbiosis (Sachs *et al.*, 2018), although almost all evidence is from SYM-type papilionoids. We suggest that the anatomical differences between legume FTs and SYMs represent different degrees to which symbionts are effectively compartmentalised, as mooted by Chomicki *et al.* (2020). The SYM-type nodule, in which the microsymbiont is released from the wall-bound IT into membrane-bound symbiosomes within the host cell, allows for a more intimate and potentially more effective and enduring symbiotic partnership where the plant has invested in the establishment of an N_2_-fixing “organelle”, which Parniske (2018) argued has only occurred in legumes and in the *Gunnera*-*Nostoc* symbiosis. In SYM-type nodules, the plant host assumes greater control of the microsymbiont and supplies more of the components required for bacteroid metabolism and N_2_-fixation (Hakoyama *et al.*, 2009; Udvardi & Poole, 2013). This more intimate endosymbiosis reaches its pinnacle in the Inverted Repeat Lacking Clade of Papilionoideae, in which swollen endo-reduplicated bacteroids that have lost their capacity for free-living growth, but which are highly efficient at fixing N, are prevalent (Oono *et al.*, 2009; Ardley & Sprent, 2021). The bacteroid in the FT, although also surrounded by a membrane analogous to the SYM membrane, remains surrounded by a cell wall, albeit a thin one which contains little pectin, which means that it is extracellular i.e. in the apoplast. Thus, while FTs represent a modest degree of compartmentalisation, at least in *Parasponia* there is evidence to suggest that FTs are not effective in controlling growth of inefficient rhizobial strains (Op den Camp *et al.*, 2012). These differing degrees of compartmentalisation provide a compelling reason why SYM-type nodulators are less likely to be affected by cheating or infiltration by inefficient microsymbionts compared to FT-type nodulator hosts, which remain in a ‘looser’ relationship with their symbionts.

There appear to be few documented examples of such extensive evolutionary losses of a key plant functional trait as those observed here for losses of nodulation. However, it is perhaps notable that one other trait that has also been repeatedly lost across mimosoid legumes, the occurrence of extrafloral nectaries (EFNs) on the leaves (Marazzi *et al.*, 2019), is also related to another plant mutualism whereby EFNs indirectly mediate ecologically important plant defence mechanisms against herbivores by attracting ants, suggesting that mutualisms are perhaps especially vulnerable to evolutionary loss.

It might be assumed that the wall of the FT would also present an additional barrier in terms of nodule O_2_ relations and host-symbiont nutrient exchange. However, the FT wall is thin, comprised mainly of cellulose, hemicelluloses, and AGP, but with reduced levels of homogalacturonan (HG) pectin components compared to the thicker and stiffer IT wall (this study), and hence presumably relatively permeable to C4-dicarboxylates and ammonia (Brewin, 2004). The presence of abundant leghaemoglobin (Lb) suggests that rhizobial FT nodules are not obviously different from SYM-type nodules in their O_2_ exchange. We propose that the role of the thin-walled FT appears to be protective i.e. it prevents the undifferentiated bacteroids going “rogue” and proliferating within the host cell, but in parallel, prevents the plant from identifying them as pathogens and attacking them. In all other respects, FT-type nodules are similar to SYM-types in still possessing a symbiosome membrane surmounting the FT which is the real point of nutrient exchange between the two partners. In short, the FT represents a kind of compromise in which the symbiosis functions well enough to benefit both partners, but in which neither partner is fully committed to.

An additional advantage of SYM-type nodules may be that it is easier for hosts to select symbionts better adapted to specific edaphic conditions, or which afford more efficient N_2_-fixation. This is in line with the greater diversity of Alpha- and Beta-rhizobial types in SYM-type nodules and the wider geographic distribution and environmental span of SYM-type nodulating legumes (Sprent *et al.*, 2017; Ardley & Sprent, 2021), compared to FT-type nodules whose microsymbionts appear to be largely limited to *Bradyrhizobium* (Fonseca *et al.*, 2012; Parker, 2015; Ardley & Sprent, 2021) and which are mostly confined to the tropics and subtropics where bradyrhizobia are dominant and widespread (Parker, 2015; Meng *et al.*, 2019).

## Conclusions

The evolution of the symbiosome in species-rich nodulating legume lineages offers a compelling explanation for the well-known but poorly understood highly uneven distribution of nodulating species richness across the N_2_-fixing clade. While nodulation has been suggested as a possible key innovation underpinning the evolutionary success of legumes, our results suggest that it was adoption of SYM-type nodules and the innovation of the symbiosome that underpinned the stabilization of N_2_-fixation and potentially contributed to massive diversification of species within Caesalpinioideae and Papilionoideae, the two most diverse and geographically widespread subfamilies of legumes. Furthermore, the greater propensity of the FT-type nodule to be secondarily lost and for SYM-type lineages to persist and diversify provides a potent example of the long-term evolutionary benefits and outcomes of stricter compartmentalisation in symbiotic cooperation (Chomicki *et al.*, 2020).

We show that the grade of caesalpinioid lineages subtending the Mimosoid clade is a hotspot of evolutionary transitions between phylogenetically intermingled nodulating and non-nodulating lineages (Fig. 1), including two independent transitions from FT- to SYM-type nodules as well as numerous losses of nodulation. The phylogeny and detailed evolutionary trajectories of nodulation and nodule anatomies presented here provide a robust framework for comparative genomic analyses of FT and SYM nodulating and non-nodulating lineages across Caesalpinioideae.

## Acknowledgements

We thank Gwilym Lewis for help with identification of plants, the Millennium Seedbank for supplying seeds, and colleagues who have helped sample material used in this study. This work is supported by the research project Engineering Nitrogen Symbiosis for Africa (ENSA), which is funded by a grant to the University of Cambridge by the Bill & Melinda Gates Foundation and the Foreign, Commonwealth & Development Office (FCDO), the Swiss National Science Foundation (grants 310003A_156140 and 31003A_182453/1 to CEH), CNPq grant 312125/2020-8 to SMdF, and Fundação de Amparo à Pesquisa do Estado da Bahia (FAPESB, grant APP0037/2016 to DC).

## Author Contributions

EJMK, JJR, CEH & EJK designed the study and interpreted the results, JJR carried out the phylogenetic analysis, SMF, EKJ, EG, KW, and YP performed anatomical analyses, SMF, EKJ, EG, DC, GKDA, JA (Ghana), NT, HSG, YP, MM, NT, PS, HL, and CZ sampled nodules in the field, SMF, EJMK, JJR, EKJ, JA (Australia), DC, EG, YP, JIS & CEH wrote the paper.

## Supporting Information

**Fig. S1** Symbiosomes are standard in nodules of caesalpinioids from the Mimosoid clade. Light (a, c, e, g, k) and transmission electron microscope (TEM) (b, d, f, h, l) images of sections of nodules from various mimosoid nodules. **a,** *Entada polystachya* nodule longitudinal profile illustrating the zonation typical of an indeterminate nodule (m = meristem, iz = invasion zone, nf = nitrogen fixing zone). Bar = 50 µm. **b,** TEM of a *E. polystachya* nf zone cell with its N-fixing bacteroids (b) contained in symbiosomes with distinct membranes (arrows) separating them from the host cytoplasm (c). Bar = 1 µm. **c,** *Enterolobium cyclocarpum* nodule longitudinal profile illustrating the zonation typical of an indeterminate nodule (m = meristem, nf = nitrogen fixing zone). Bar = 200 µm. **d,** TEM of an *E. cyclocarpum* nf zone cell with its bacteroids (b) enclosed in symbiosomes (*) that are separated from the host cytoplasm (c) by the symbiosome membrane (arrows). An IT is also present in the cell; its wall (w) is immunogold labelled with 10 nm gold particles linked to JIM5 (arrowheads). Bar = 1 µm. **e,** *Lachesiodendron viridiflorum* nodule longitudinal profile illustrating the zonation typical of an indeterminate nodule (m = meristem, nf = nitrogen fixing zone). Bar = 200 µm. **f,** TEM of an *L. viridiflorum* nf zone cell with its bacteroids (b) enclosed in symbiosomes (*) that are separated from the host cytoplasm (c) by the symbiosome membrane (arrows). Bar = 500 nm. **g,** high magnification view of the nf zone of a *Xylia xylocarpa* nodule showing large bacteroid-containing cells (b) surrounded by smaller and more numerous uninfected cells (u). Bar = 25 µm. **h,** TEM of an *X. xylocarpa* nf zone cell with its bacteroids (b) enclosed in symbiosomes that are separated from the host cytoplasm (c) by the symbiosome membrane (arrows). Bar = 500 nm.

**Fig. S2** Fixation threads (FTs) are standard in non-mimosoid grade Caesalpinioid nodules. Light (a, c, e, g, k) and transmission electron microscope (b, d, f, h, l) images of sections of nodules from *Moldenhawera* spp. (a – f), and various other caesalpinioid nodules (g – l). **a,** whole *M. blanchetiana* var. *multijuga* nodule longitudinal profile illustrating the zonation typical of an indeterminate nodule (m = meristem, iz = invasion zone, nf = nitrogen fixing zone). Bar = 200 µm. **b,** higher magnification view of the nf zone of an *M. multijuga* nodule showing large bacteroid-containing cells (b) surrounded by smaller and more numerous uninfected cells (u). Bar = 25 µm. **c,** *M. floribunda* iz cell with an infection thread (IT) containing a bacterium (b) within a strand of cytoplasm (c); the wall of the IT is densely immunogold labelled with 10 nm gold particles linked to JIM5 (arrows). v = vacuole. Bar = 1 µm. **d,** *M. blanchetiana* var. *multijuga* bacteroids (b) within FTs adjacent to the host cell wall (w) which is immunogold labelled with 10 nm gold particles linked to JIM5 (arrows); note that the FT walls are almost completely unlabelled (a few solitary gold particles are indicated by arrowheads). c = host cytoplasm. Bar = 500 nm. **e,** young nf cell in a *M. floribunda* nodule in which the few bacteria-containing FTs within it are still confined to strands of cytoplasm (c) adjacent to the nucleus (n). Note the strand of plasma membrane (double arrowhead) associated with an FT; it appears to be derived from the nuclear membrane (arrowheads). v = vacuole. Bar = 1 µm. **f,** mature FTs in the nf of a *M. floribunda* nodule. Note the thick wall (w) of the FTs and the membranes associated with them (arrows). c = cytoplasm, m = mitochondrion. Bar = 500 nm. **g,** high magnification view of the nf zone of a *Jacqueshuberia purpurea* nodule showing large bacteroid-containing cells (b) surrounded by smaller and more numerous uninfected cells (u). Bar = 25 µm. h, TEM of a *J. purpurea* nf zone cell with its bacteroids (b) enclosed in FTs that are separated from the host cytoplasm (c) by electron-dense walls (arrows) that are almost completely unlabelled with JIM5, except for some thickened areas (arrowheads). Bar = 500 nm. **i,** high magnification view of the nf zone of a *Tachigali rugosa* nodule showing large bacteroid-containing cells (b) surrounded by smaller uninfected cells (u). Bar = 25 µm. **j,** TEM of a *T. rugosa* nf zone cell with its bacteroids (b) enclosed in FTs that are separated from the host cytoplasm (c) by cell walls that are labelled with JIM5 (arrows). v = vacuole, m = mitochondrion. Bar = 1 µm. **k,** high magnification view of the nf zone of a *Campsiandra comosa* nodule showing large bacteroid-containing cells (b) surrounded by smaller uninfected cells (u). Bar = 25 µm. **l,** TEM of a *C. comosa* nf zone cell with its bacteroids (b) enclosed in FTs that are separated from the host cytoplasm (c) by cell walls that are labelled with JIM5 (arrows). Bar = 500 nm.

**Fig. S3** Evolutionary trajectory of nodulation and nodule type when taxa with missing data have been assigned equal weight to nodulation and non-nodulation states. Methods and legend otherwise as for Figure 1.

**Table S1.** Caesalpinioideae and outgroup taxa used in the time-calibrated phylogeny depicting evolutionary trajectories of nodulation and nodule type (Fig. 1, Fig. S3). The nodulation status of each genus is recorded as confirmed nodulated (Green), Non-nodulated (blue) or Unknown (orange). Nodule anatomy is recorded (if known) with regard to the presence of fixation threads (FTs) or symbiosomes.

**Table S2.** Occurrence of fixation threads (FTs) and/or symbiosomes in nodules from Caesalpinioideae (Mim, belongs to the Mimosoid clade) extracted from the literature and from the unpublished observations of the authors. Details of the references are in Notes S1.

**Table S3.** Type, location, and age of transitions in nodulation status as depicted in Figure 1. Transitions are inferred to have occurred on the stem of each taxon or clade listed in the Location column.

**Notes S1.** References for Table S2.

## Notes

### Competing Interest Statement

The authors have declared no competing interest.

## References

Ardley J, Sprent J. 2021. Evolution and biogeography of actinorhizal plants and legumes: A comparison. Journal of Ecology 109: 1098–1121.

Batterman SA, Hedin LO, Van Breugel M, Ransijn J, Craven DJ, Hall JS. 2013. Key role of symbiotic dinitrogen fixation in tropical forest secondary succession. Nature 502: 224–227.

Brewin NJ. 2004. Plant cell wall remodelling in the rhizobium-legume symbiosis. Critical Reviews in Plant Sciences 23: 293–316.

Bruneau A, Mercure M, Lewis GP, Herendeen PS. 2008. Phylogenetic patterns and diversification in the caesalpinioid legumes. Botany 86: 697–718.

Cardoso D, Pennington RT, de Queiroz LP, Boatwright JS, Van Wyk BE, Wojciechowski MF, Lavin M. 2013. Reconstructing the deep-branching relationships of the papilionoid legumes. South African Journal of Botany 89: 58–75.

Cardoso D, De Queiroz LP, Toby Pennington R, De Lima HC, Fonty É, Wojciechowski MF, Lavin M. 2012. Revisiting the phylogeny of papilionoid legumes: New insights from comprehensively sampled early-branching lineages. American Journal of Botany 99: 1991–2013.

Chen H, Markham J. 2021. Ancient CO2 levels favor nitrogen fixing plants over a broader range of soil N compared to present. Scientific Reports 11: 1–6.

Chomicki G, Werner GDA, West SA, Kiers ET. 2020. Compartmentalization drives the evolution of symbiotic cooperation. Philosophical Transactions of the Royal Society B 375: 20190602.

Doyle JJ. 2011. Phylogenetic perspectives on the origins of nodulation. Molecular Plant-Microbe Interactions 24: 1289–1295.

Doyle JJ. 2016. Chasing unicorns: Nodulation origins and the paradox of novelty. American Journal of Botany 103: 1865–1868.

Drummond AJ, Rambaut A. 2007. BEAST: Bayesian evolutionary analysis by sampling trees. BMC Evolutionary Biology 7: 1–8.

Edwards EJ. 2019. Evolutionary trajectories, accessibility and other metaphors: the case of C4 and CAM photosynthesis. New Phytologist 223: 1742–1755.

Epihov DZ, Batterman SA, Hedin LO, Leake JR, Smith LM, Beerling DJ. 2017. N2-fixing tropical legume evolution: A contributor to enhanced weathering through the Cenozoic? Proceedings of the Royal Society B: Biological Sciences 284: 20170370.

de Faria SM, McInroy SG, Sprent JI. 1987. The occurrence of infected cells, with persistent infection threads, in legume root nodules. Canadian Journal of Botany 65: 553–558.

de Faria SM, Sutherland JM, Sprent JI. 1986. A new type of infected cell in root nodules of Andira spp. (Leguminosae). Plant Science 45: 143–147.

Fonseca MB, Peix A, de Faria SM, Mateos PF, Rivera LP, Simões-Araujo JL, França MGC, dos Santos Isaias RM, Cruz C, Velázquez E, et al. 2012. Nodulation in Dimorphandra wilsonii Rizz. (Caesalpinioideae), a Threatened Species Native to the Brazilian Cerrado. PLoS ONE 7: 1–16.

Frank SA. 1996. Host — symbiont conflict over the mixing of symbiotic lineages. Proceedings of Royal Society of London B 263: 339–344.

Gagnon E, Bruneau A, Hughes CE, de Queiroz L, Lewis GP. 2016. A new generic system for the pantropical Caesalpinia group (Leguminosae). PhytoKeys 71: 1–160.

Griesmann M, Chang Y, Liu X, Song Y, Haberer G, Crook MB, Billault-Penneteau B, Lauressergues D, Keller J, Imanishi L, et al. 2018. Phylogenomics reveals multiple losses of nitrogen-fixing root nodule symbiosis. Science 361: eaat1743.

Hakoyama T, Niimi K, Watanabe H, Tabata R, Matsubara J, Sato S, Nakamura Y, Tabata S, Jichun L, Matsumoto T, et al. 2009. Host plant genome overcomes the lack of a bacterial gene for symbiotic nitrogen fixation. Nature 462: 514–517.

Hoffman BM, Lukoyanov D, Yang ZY, Dean DR, Seefeldt LC. 2014. Mechanism of nitrogen fixation by nitrogenase: The next stage. Chemical Reviews 114: 4041–4062.

Kiers ET, Rousseau RA, West SA, Denison RF. 2003. Host sanctions and the legume – rhizobium mutualism. Nature 425: 78–81.

Koenen EJM, Kidner CA, de Souza ÉR, Simon MF, Iganci JR V., Nicholls JA, Brown GK, de Queiroz LP, Luckow MA, Lewis GP, et al. 2020a. Hybrid capture of 964 nuclear genes resolves evolutionary relationships in the mimosoid legumes and reveals the polytomous origins of a large pantropical radiation. American Journal of Botany 107: 1710–1735.

Koenen EJM, Ojeda DI, Steeves R, Migliore J, Bakker FT, Wieringa JJ, Kidner C, Hardy OJ, Pennington RT, Bruneau A, et al. 2020b. Large‐scale genomic sequence data resolve the deepest divergences in the legume phylogeny and support a near‐simultaneous evolutionary origin of all six subfamilies. New Phytologist 225: 1355–1369.

De La Peña TC, Fedorova E, Pueyo JJ, Mercedes Lucas M. 2018. The symbiosome: Legume and rhizobia co-evolution toward a nitrogen-fixing organelle? Frontiers in Plant Science 8: 2229.

Lancelle SA, Torrey JG. 1985. Early development of Rhizobium-induced root nodules of Parasponia rigida. I. Infection and early nodule initiation. Canadian Journal of Botany 63: 25–35.

Ledermann R, Schulte CCM, Poole PS. 2021. How rhizobia adapt to the nodule environment. Journal of Bacteriology 203: e00539–20.

LPWG. 2017. A new subfamily classification of the Leguminosae based on a taxonomically comprehensive phylogeny – The Legume Phylogeny Working Group (LPWG). Taxon 66: 44–77.

Manzanilla V, Bruneau A. 2012. Phylogeny reconstruction in the Caesalpinieae grade (Leguminosae) based on duplicated copies of the sucrose synthase gene and plastid markers. Molecular Phylogenetics and Evolution 65: 149–162.

Marazzi B, Ané C, Simon MF, Delgado-Salinas A, Luckow M, Sanderson MJ. 2012. Locating evolutionary precursors on a phylogenetic tree. Evolution 66: 3918–3930.

Marazzi B, Gonzalez AM, Delgado-Salinas A, Luckow MA, Ringelberg JJ, Hughes CE. 2019. Extrafloral nectaries in Leguminosae: phylogenetic distribution, morphological diversity and evolution. Australian Systematic Botany 32: 409–458.

McKey D. 1994. Legumes and nitrogen: the evolutionary ecology of a nitrogen-demanding lifestyle. In: Advances in Legume Systematics 5: The Nitrogen Factor. 211–228.

Meng H, Zhou Z, Wu R, Wang Y, Gu JD. 2019. Diazotrophic microbial community and abundance in acidic subtropical natural and re-vegetated forest soils revealed by high-throughput sequencing of nifH gene. Applied Microbiology and Biotechnology 103: 995–1005.

Mergaert P, Uchiumi T, Alunni B, Evanno G, Cheron A, Catrice O, Mausset A-E, Barloy-Hubler F, Galibert F, Kondorosi A, et al. 2006. Eukaryotic control on bacterial cell cycle and differentiation in the Rhizobium – legume symbiosis. Proceedings of the National Academy of Sciences 103: 5230–5235.

Naisbitt T, James EK, Sprent JI. 1992. The evolutionary significance of the legume genus Chamaecrista, as determined by nodule structure. New Phytologist 122: 487–492.

Oldroyd GED. 2013. Speak, friend, and enter: Signalling systems that promote beneficial symbiotic associations in plants. Nature Reviews Microbiology 11: 252–263.

Oldroyd GED, Downie JA. 2008. Coordinating nodule morphogenesis with rhizobial infection in legumes. Annual Review of Plant Biology 59: 519–546.

Oono R, Denison RF, Kiers ET. 2009. Controlling the reproductive fate of rhizobia: how universal are legume sanctions? New Phytologist 183: 967–979.

Op den Camp RHM, Polone E, Fedorova E, Roelofsen W, Squartini A, Den Camp HJMO, Bisseling T, Geurts R. 2012. Nonlegume Parasponia andersonii deploys a broad rhizobium host range strategy resulting in largely variable symbiotic effectiveness. Molecular Plant-Microbe Interactions 25: 954–963.

Parker MA. 2015. The Spread of Bradyrhizobium Lineages Across Host Legume Clades: from Abarema to Zygia. Microbial Ecology 69: 630–640.

Parniske M. 2018. Uptake of bacteria into living plant cells, the unifying and distinct feature of the nitrogen-fixing root nodule symbiosis. Current Opinion in Plant Biology 44: 164–174.

Pawlowski K, Demchenko KN. 2012. The diversity of actinorhizal symbiosis. Protoplasma 249: 967–979.

Peoples MB, Herridge DF, Ladha JK. 1995. Biological nitrogen fixation: An efficient source of nitrogen for sustainable agricultural production? Plant and Soil 174: 3–28.

Revell LJ. 2012. phytools: An R package for phylogenetic comparative biology (and other things). Methods in Ecology and Evolution 3: 217–223.

Ribeiro PG, Luckow M, Lewis GP, Simon MF, Cardoso D, de Souza ÉR, Silva APC, Jesus MC, dos Santos FAR, Azevedo V, et al. 2018. Lachesiodendron, a new monospecific genus segregated from Piptadenia (Leguminosae: Caesalpinioideae: Mimosoid clade): Evidence from morphology and molecules. Taxon 67: 37–54.

Rogers A, Ainsworth EA, Leakey ADB. 2009. Will elevated carbon dioxide concentration amplify the benefits of nitrogen fixation in legumes? Plant Physiology 151: 1009–1016.

Rubio MC, Becana M, Kanematsu S, Ushimaru T, James EK. 2009. Immunolocalization of antioxidant enzymes in high-pressure frozen root and stem nodules of Sesbania rostrata. New Phytologist 183: 395–407.

Sachs JL, Quides KW, Wendlandt CE. 2018. Legumes versus rhizobia: a model for ongoing conflict in symbiosis. New Phytologist 219: 1199–1206.

dos Santos JMF, Casaes Alves PA, Silva VC, Kruschewsky Rhem MF, James EK, Gross E. 2017. Diverse genotypes of Bradyrhizobium nodulate herbaceous Chamaecrista (Moench) (Fabaceae, Caesalpinioideae) species in Brazil. Systematic and Applied Microbiology 40: 69–79.

Shen D, Bisseling T. 2020. The evolutionary aspects of legume nitrogen–fixing nodule symbiosis. In: Symbiosis: Cellular, Molecular, Medical and Evolutionary Aspects. Cham, Switzerland: Springer, 387–408.

Simon MF, Pastore JFB, Souza AF, Borges LM, Scalon VR, Ribeiro PG, Santos-Silva J, Souza VC, Queiroz LP. 2016. Molecular Phylogeny of Stryphnodendron (Mimosoideae, Leguminosae) and Generic Delimitations in the Piptadenia Group. International Journal of Plant Sciences 177: 44–59.

Smith CA, Skvirsky RC, Hirsch AM. 1986. Histochemical evidence for the presence of a suberinlike compound in Rhizobium -induced nodules of the nonlegume Parasponia rigida. Canadian Journal of Botany 64: 1474–1483.

Soltis DE, Soltis PS, Morgan DR, Swensen SM, Mullin BC, Dowd JM, Martin PG. 1995. Chloroplast gene sequence data suggest a single origin of the predisposition for symbiotic nitrogen fixation in angiosperms. Proceedings of the National Academy of Sciences of the United States of America 92: 2647–2651.

Sprent JI. 2001. Nodulation in Legumes. Richmond, United Kingdom: Royal Botanic Gardens Kew.

Sprent JI. 2009. Legume Nodulation: A Global Perspective. Oxford, United Kingdom: Wiley-Blackwell.

Sprent JI, Ardley JK, James EK. 2013. From North to South: A latitudinal look at legume nodulation processes. South African Journal of Botany 89: 31–41.

Sprent JI, Ardley J, James EK. 2017. Biogeography of nodulated legumes and their nitrogen-fixing symbionts. New Phytologist 215: 40–56.

Taylor BN, Menge DNL. 2018. Light regulates tropical symbiotic nitrogen fixation more strongly than soil nitrogen. Nature Plants 4: 655–661.

Trinick MJ. 1980. Effects of Oxygen, Temperature and Other Factors on the Reduction of Acetylene by Root Nodules Formed by Rhizobium on Parasponia andersonii Planch. New Phytologist 86: 27–38.

Tsyganova A V., Brewin NJ, Tsyganov VE. 2021. Structure and development of the legume-rhizobial symbiotic interface in infection threads. Cells 10: 1050.

Udvardi M, Poole PS. 2013. Transport and metabolism in legume-rhizobia symbioses. Annual Review of Plant Biology 64: 781–805.

VandenBosch KA, Bradley DJ, Knox JP, Perotto S, Butcher GW, Brewin NJ. 1989. Common components of the infection thread matrix and the intercellular space identified by immunocytochemical analysis of pea nodules and uninfected roots. The EMBO Journal 8: 335–341.

van Velzen R, Doyle JJ, Geurts R. 2018a. A Resurrected Scenario: Single Gain and Massive Loss of Nitrogen-Fixing Nodulation. Trends in Plant Science 24: 49–57.

van Velzen R, Holmer R, Bu F, Rutten L, van Zeijl A, Liu W, Santuari L, Cao Q, Sharma T, Shen D, et al. 2018b. Comparative genomics of the nonlegume Parasponia reveals insights into evolution of nitrogen-fixing rhizobium symbioses. Proceedings of the National Academy of Sciences of the United States of America 115: E4700–E4709.

Vitousek PM, Menge DNL, Reed SC, Cleveland CC. 2013. Biological nitrogen fixation: Rates, patterns and ecological controls in terrestrial ecosystems. Philosophical Transactions of the Royal Society B: Biological Sciences 368: 20130119.

Werner GDA, Cornwell WK, Sprent JI, Kattge J, Kiers ET. 2014. A single evolutionary innovation drives the deep evolution of symbiotic N2-fixation in angiosperms. Nature Communications 5: 1–9.

White J, Prell J, James EK, Poole P. 2007. Nutrient sharing between symbionts. Plant Physiology 144: 604–614.

Yang S, Wang Q, Fedorova E, Liu J, Qin Q, Zheng Q, Price PA, Pan H, Wang D, Griffitts JS, et al. 2017. Microsymbiont discrimination mediated by a host-secreted peptide in Medicago truncatula. Proceedings of the National Academy of Sciences 114: 6848–6853.

Zhang C, Rabiee M, Sayyari E, Mirarab S. 2018. ASTRAL-III: Polynomial time species tree reconstruction from partially resolved gene trees. BMC Bioinformatics 19: 15–30.

Zhao Y, Zhang R, Jiang K, Qi J, Hu Y, Guo J, Zhu R, Zhang T, Egan AN, Yi T-S, et al. 2021. Nuclear Phylotranscriptomics/Phylogenomics Support Numerous Polyploidization Events and Hypotheses for the Evolution of Rhizobial Nitrogen-Fixing Symbiosis in Fabaceae. Molecular Plant.

## References for Table S2

Baird LM, Virginia RA, Webster BD (1985) Development of Root Nodules in a Woody Legume, *Prosopis glandulosa* Torr. Botanical Gazette 146: 39–43.

Beukes CW, Boshoff FS, Phalane FL, Hassen AI, le Roux MM, Stepkowski T, Venter SN, Steenkamp ET (2019) Both alpha and beta-rhizobia occupy the root nodules of *Vachellia karroo* in South Africa. Front Microbiol 10:1195.

Bontemps, C., Rogel, M.A., Wiechmann, A., Mussabekova, A., Moody, S., Simon, M.F., Moulin, L., Elliott, G.N., Lacercat-Didier, L., Dasilva, C., Grether, R., Camargo-Ricalde, S.L., Chen, W., Sprent, J.I., Martínez-Romero, E., Young, J.P.W., and James, E.K. (2016) Endemic *Mimosa* species from Mexico prefer alphaproteobacterial rhizobial symbionts. New Phytologist 209, 319–333.

Bournaud C., de Faria, S.M., dos Santos, J.M.F., Tisseyre, P., Silva, M., Chaintreuil, C., Gross, E., James, E.K., Prin, Y., and Moulin, L. (2013) *Burkholderia* species are the most common and preferred nodulating symbionts of the Piptadenia Group (tribe Mimoseae). PLoS ONE 8, 63476.

Bournaud C., James, E.K., de Faria, S.M., Lebrun, M., Melkonian, R., Duponnois, R., Tisseyre, P., Moulin, L. and Prin, Y. (2018) Interdependency of efficient nodulation and arbuscular mycorrhization in *Piptadenia gonoacantha,* a Brazilian legume tree. Plant, Cell and Environment 41, 2008 – 2020.

Canosa GA, de Faria SM, de Moraes LFD (2012) Leguminosas florestais da Mata Atlántica brasileira fixadoras de nitrogênio atmosférico. Embrapa Communicado Técnico 144 ISSN 1517-8862

Chen, W-M., James, E.K., Prescott, A.R., Kierans, M., and Sprent, J.I. (2003) Nodulation of *Mimosa* spp. by the β-proteobacterium *Ralstonia taiwanensis*. Molecular Plant-Microbe Interactions 16, 1051–1061.

Chen, W-M., de Faria, S.M., Straliotto, R., Pitard, R.M., Simões-Araùjo, J.L., Chou, Yi-Ju Chou, J-H., Barrios, E., Prescott, A.R., Elliott, G.N., Sprent, J.I., Young, J.P.W., and James, E.K. (2005) Proof that *Burkholderia* Forms Effective Symbioses with Legumes: a Study of Novel *Mimosa*-nodulating Strains from South America. Applied and Environmental Microbiology 71, 7461–7471.

Chen, W-M., James, E.K., Chou, J-H, Sheu, S-Y., Yang, S-Z., and Sprent, J.I. (2005) Beta-rhizobia from *Mimosa pigra*, a newly-discovered invasive plant in Taiwan. New Phytologist 168, 661–675.

Choudhary, C., Tak, N., Bissa, G., Chouhan, B., Choudhary, P., Sprent, J.I., James, E.K., and Gehlot H.S. (2020) The widely distributed legume tree *Vachellia* (*Acacia*) *nilotica* subsp. *indica* is nodulated by genetically diverse *Ensifer* strains in India. Symbiosis 80, 15–31.

Cordero I, Ruiz-Díez B, Coba de la Peña T, Balaguer L, Lucas MM, Rincón A, Pueyo JJ (2016) Rhizobial diversity, symbiotic effectiveness and structure of nodules of *Vachellia macracantha*. Soil Biol Biochem 96: 39–54. doi.org/10.1016/j.soilbio.2016.01.011

Diabate M, Munive A, Faria SM de, Ba A, Dreyfus B, Galiana A (2005) Occurrence of nodulation in unexplored leguminous trees native to the West African tropical rainforest and inoculation response of native species useful in reforestation. New Phytol 166: 231–239.

Dupuy, N (1993) Contribution à l’étude de la symbiose fixatrie d’azote entre *Acacia albida* et *Bradyrhizobium* sp. Thèse de doctorat. Université de Lille.

Elliott, G.N., Chen, W-M., Chou, J-H., Wang, H-C., Sheu, S-Y., Perin, L., Reis, V.M., Moulin, L., Simon, M.F., Bontemps, C., Sutherland, J.M., Bessi, R., de Faria, S.M., Trinick, M.J., Prescott, A.R., Sprent, J.I. and James, E.K. (2007) *Burkholderia phymatum* is a highly effective nitrogen-fixing symbiont of *Mimosa* spp. and fixes nitrogen *ex planta*. New Phytologist 173, 168–180.

Elliott, G.N., Chou, J-H., Chen, W-M., Bloemberg, G.V., Bontemps, C., Martínez-Romero, E., Velázquez, E., Young, J.P.W., Sprent, J.I. and James, E.K. (2009) *Burkholderia* spp. are the most competitive symbionts of *Mimosa*, particularly under N-limited conditions. Environmental Microbiology 11, 762–778.

Faria SM de, Lima HC de (1998) Additional studies of the nodulation status of legume species in Brazil. Plant and Soil 200, 185–192.

Faria SM de, Lima HC de (2002) Levantamento de nodulação em leguminosas arbóreas e arbustivas em áreas de influência da Mineração Rio do Norte – Porto Trombetas / PA. Embrapa Agrobiologia. Documentos 159. 32 pp. ISSN 1517-8498 http://www.infoteca.cnptia.embrapa.br/infoteca/handle/doc/624782

Faria SM de, Lima HC de, Ribeiro RD, Castilho AF, Henriques JC (2006) Nodulação em espécies leguminosas da região de Porto Trombetas, Oriximiná, Estado do Pará e seu potencial uso no reflorestamento de bacias de rejeito do lavado de bauxita. Embrapa Agrobiologia. Documentos 209. 14 pp. ISSN 1517-8498 http://www.infoteca.cnptia.embrapa.br/infoteca/handle/doc/628849

Faria SM de, Diedhiou AG, Lima HC de, Ribeiro RD, G, Castilho AF, Galiana A, Henriques JC (2010) Evaluating the nodulation status of leguminous species from the Amazonian forest of Brazil. J Exp Bot 61: 3119–3127.

Faria SM de, Moraes, LFD de,Lima HC de, Ribeiro RD, Mattos, CMJ, Rodrigues TM, Castilho AF, Canosa GA, Silva MAP (2011) Composição florística de leguminosas com potencial para fixação biológica de nitrogênio em áreas de vegetação de Canga (savana metalófita) do entorno do complexo minerador de Carajás. Embrapa Agrobiologia. Communicado Técnico 140. 20 pp. ISSN 1517-8862 http://www.infoteca.cnptia.embrapa.br/infoteca/handle/doc/921085

Faria SM de, McInroy SG, Sprent JI (1987) The occurrence of infected cells, with persistent infection threads, in legume root nodules. Can J Bot 65: 553–558.

Fonseca, M.B., Peix, A., de Faria, S.M., Mateos, P.F., Rivera, L.P., Simões-Araujo, J.L., Costa França, M.G., dos Santos Isaias, R.M., Cruz, C., Velázquez, E., Scotti, M.R., Sprent, J.I. and James, E.K. (2012) Nodulation in *Dimorphandra wilsonii* Rizz. (Caesalpinioideae), a threatened species native to the Brazilian Cerrado. PLoS ONE 7, e49520

Gehlot, H.S., Tak, N., Kaushik, M., Mitra, S., Chen, W-M., Poweleit, N., Panwar, D., Poonar, N., Parihar, R., Tak, A., Sankhla, I.S., Ojha, A., Rao, S.R., Simon, M.F., dos Reis Junior, F.B., Perigolo, N., Tripathi, A., Sprent, J.I., Young, J.P.W., James, E.K., and Gyaneshwar P. (2013) An invasive *Mimosa* in India does not adopt the symbionts of its native relatives. Annals of Botany 112: 179–196.

Gross E, Cordeiro L, Caetano FH (2002) Nodule ultrastructure and initial growth of *Anadenanthera peregrina* (L.) Speg. var. *falcata* (Benth.) Altschul plants infected with rhizobia. Ann Bot 90:175–183. doi:10.1093/aob/mcf184

Gyaneshwar, P., Hirsch, A.M., Moulin, L., Chen, W-M., Elliott, G.N., Bontemps, C., Estrada-de los Santos, P., Gross, E., dos Reis Junior, F.B., Sprent, J.I., Young, J.P.W. and James, E.K. (2011) Legume-nodulating betaproteobacteria: diversity, host range and future prospects. Molecular Plant-Microbe Interactions 24, 1276–1288.

James, E.K., Sprent, J.I., Sutherland, J.M., McInroy, S.G. and Minchin, F.R. (1992) The structure of nitrogen fixing nodules on the aquatic mimosoid legume *Neptunia plena*. Annals of Botany 69, 173–180.

Laste KCD, Gonçalves FS, Faria SM de (2008) Estirpes de rizóbio eficientes na fixação biológica de nitrogênio para leguminosas com potencial de uso na recuperação de áreas mineradas. Embrapa Agrobiologia. Comunicado Técnico, 115, 8 pp. ISSN 1517-8862 http://www.infoteca.cnptia.embrapa.br/infoteca/handle/doc/629821

Moulin L., Klonowska, A., Bournaud, C., Booth, K., Vriezen, J.A.C., Melkonian, R., James, E.K., Young, J.P.W., Bena, G., Hauser, L., Land, M., Kyrpides, N., Bruce, D., Chain, P., Copeland, A., Pitluck, S., Woyke, T., Lizotte-Waniewski, M., Bristow, J., Riley, M. (2014) Complete Genome sequence of *Burkholderia phymatum* STM815^T^, a broad host range and efficient nitrogen-fixing symbiont of *Mimosa* species. Standards in Genomic Sciences 9, 763–774.

Naisbitt, T., James E.K. and Sprent J.I. (1992) The evolutionary significance of the legume genus *Chamaecrista*, as determined by nodule structure. New Phytologist 122, 487–492.

Perrineau MM, Galiana A, de Faria SM, Bena G, Duponnois R, Reddell R, Prin Y (2012) Monoxenic nodulation process of *Acacia mangium* (Mimosoideae, Phyllodineae) by *Bradyrhizobium* sp. Symbiosis 56:87–95. doi:10.1007/s13199-012-0163-5

Platero, R., James, E.K., Rios, C., Iriarte, A., Sandes, L., Zabaleta, M., Battistoni, F., and Fabiano, E. (2016) Novel *Cupriavidus* strains isolated from root nodules of native Uruguayan *Mimosa* species. Applied and Environmental Microbiology 82, 3150 –3164.

Rathi, S., Tak, N., Bissa, G., Chouhan, B., Ojha, A., Adhikari, D., Barik, S.K., Satyawada, R.R., Sprent, J.I., James, E.K., and Gehlot H.S. (2018) Selection of *Bradyrhizobium* or *Ensifer* symbionts by the native Indian caesalpinioid legume *Chamaecrista pumila* depends on soil pH and other edaphic and climatic factors. FEMS Microbiology Ecology 10.1093/femsec/fiy180

dos Reis Junior, F.B., Simon, M.F., Gross, E., Boddey, R.M., Elliott, G.N., Neto, N.E., Loureiro, M.F., Queiroz, L.P., Scotti, M.R., Chen, W-M., Norén, A., Rubio, M.C., de Faria, S.M., Bontemps, C., Goi, S.R., Young, J.P.W., Sprent, J.I., and James, E.K. (2010) Nodulation and nitrogen fixation by *Mimosa* spp. in the Cerrado and Caatinga biomes of Brazil. New Phytologist 186, 934–946.

Rhem, M.F.K., Cordeiro Silva, V., Ferreira dos Santos, J.M., Zilli, J.E., James, E.K., Simon, M.F., and Gross, E. (2021) The large mimosoid genus *Inga* Mill. (tribe Ingeae, Caesalpinioideae) is nodulated by diverse *Bradyrhizobium* strains in its main centers of diversity in Brazil. Systematic and Applied Microbiology 44, 126268doi: https://doi.org/10.1016/j.syapm.2021.126268

Saad, MM, Crèvecoeur, M, Masson-Boivin, C, Perret X (2012) The Type 3 Protein Secretion System of *Cupriavidus taiwanensis* strain LMG19424 compromises symbiosis with *Leucaena leucocephala*. Appl. Environ Microbiol 78: 7476 – 7479.

Sankhla, I.S., Tak, N., Meghwal, R.R., Choudhary, S., Tak, A., Rathi, S., Sprent, J.I., James, E.K. and Gehlot, H.S. (2017) Molecular characterization of nitrogen fixing microsymbionts from root nodules of *Vachellia* (*Acacia*) *jacquemontii*, a native legume from the Thar Desert of India. Plant and Soil 410, 21–40

dos Santos, J.M.F., Casaes, P.A., Silva, V.C., Rhem, M.F.K., James, E.K., and Gross, E. (2017) Diverse genotypes of *Bradyrhizobium* nodulate herbaceous *Chamaecrista* (Moench) (Fabaceae, Caesalpinioideae) species in Brazil. Systematic and Applied Microbiology 40, 69–79.

Silva, V.C., Casaes, P.A., Rhem, M.F.K., dos Santos, J.M.F., James, E.K., and Gross, E. (2018) Brazilian species of *Calliandra* Benth. (tribe Ingeae) are nodulated by diverse strains of *Paraburkholderia*. Systematic and Applied Microbiology 41, 241–250.

Sprent JI. 2001. Nodulation in legumes. London, UK: Royal Botanic Gardens, Kew.

Subba Rao NS, Mateos PF, Baker D, Pankratz HS, Palma J, Dazzo FB, Sprent JI. 1995. The unique root-nodule symbiosis between *Rhizobium* and the aquatic legume, *Neptunia natans* (L. f.) Druce. Planta 196: 311–320.

Zilli, J.E., Pereira de Moraes Carvalho, C., Vieira de Matos Macedo, A., de Barros Soares, L.H., Gross, E., James, E.K., Simon, M.F., and de Faria S.M. (2021) Nodulation of the neotropical legume genus *Calliandra* by Alpha or Betaproteobacterial symbionts is dependent on the biogeographical origins of the host species. Brazilian Journal of Microbiology https://doi.org/10.1007/s42770-021-00570-8

