## Supplementary figures and images for "The innovation of the symbiosome has enhanced the evolutionary stability of nitrogen fixation in legumes"

### Figure S1

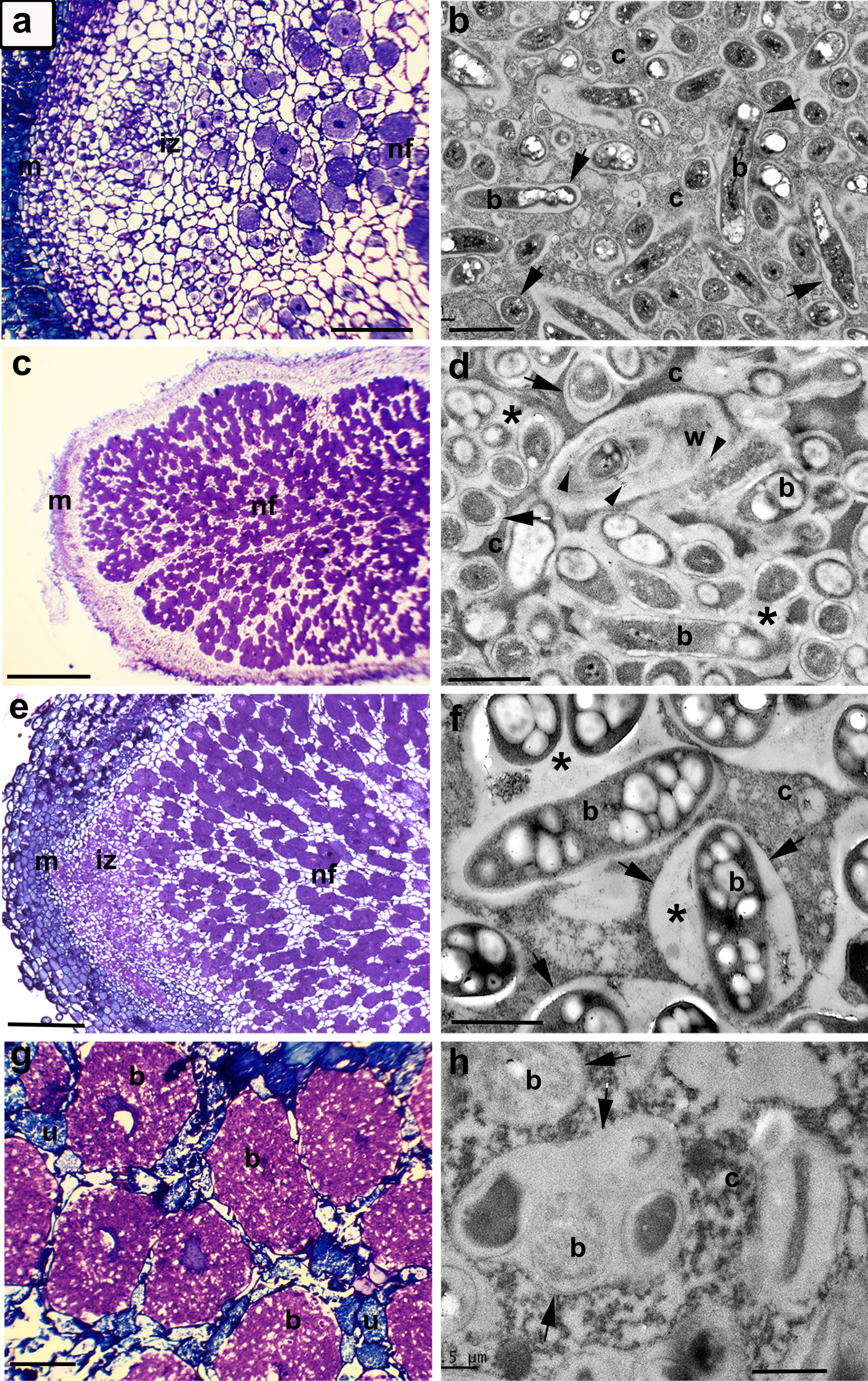

### Figure S2

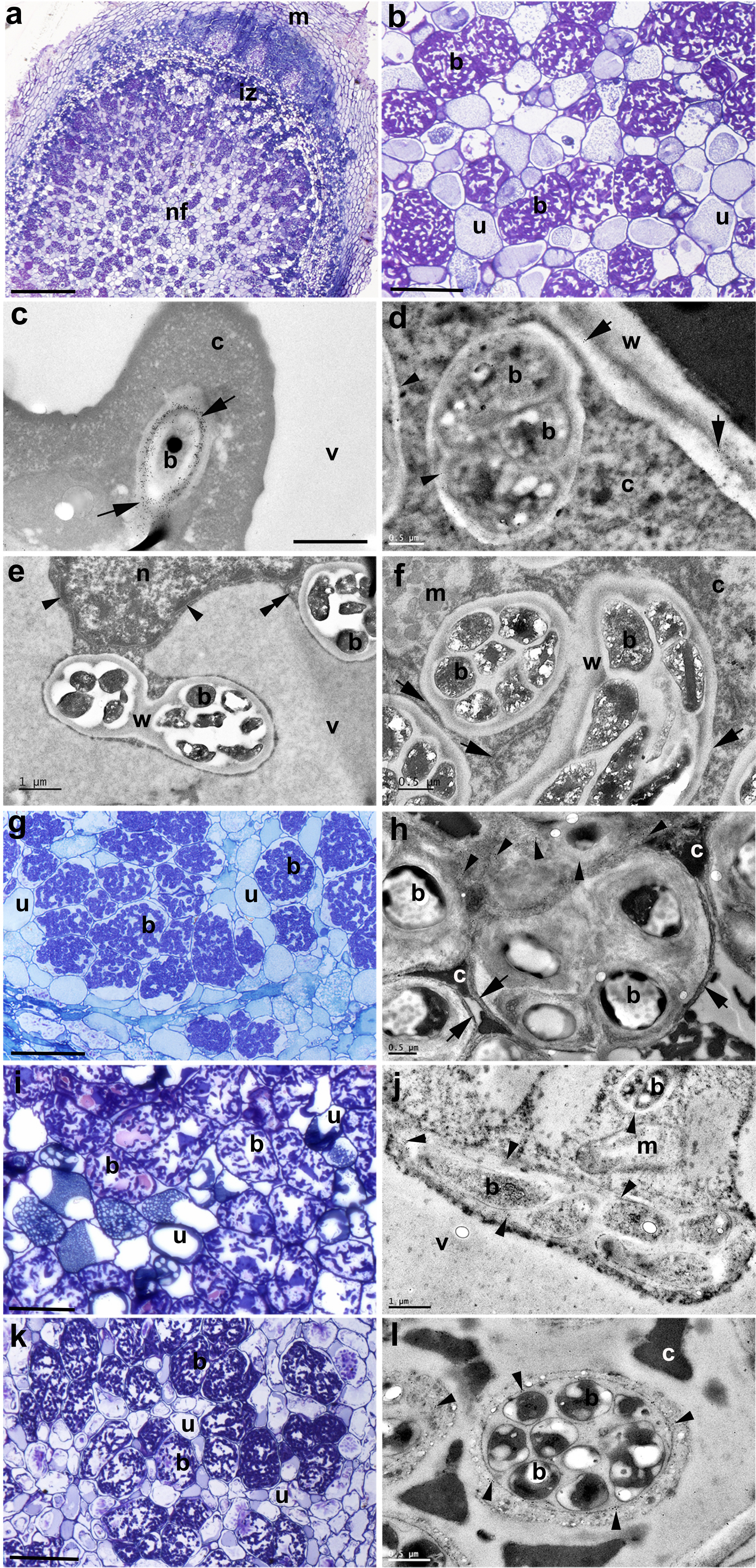

### Figure S3

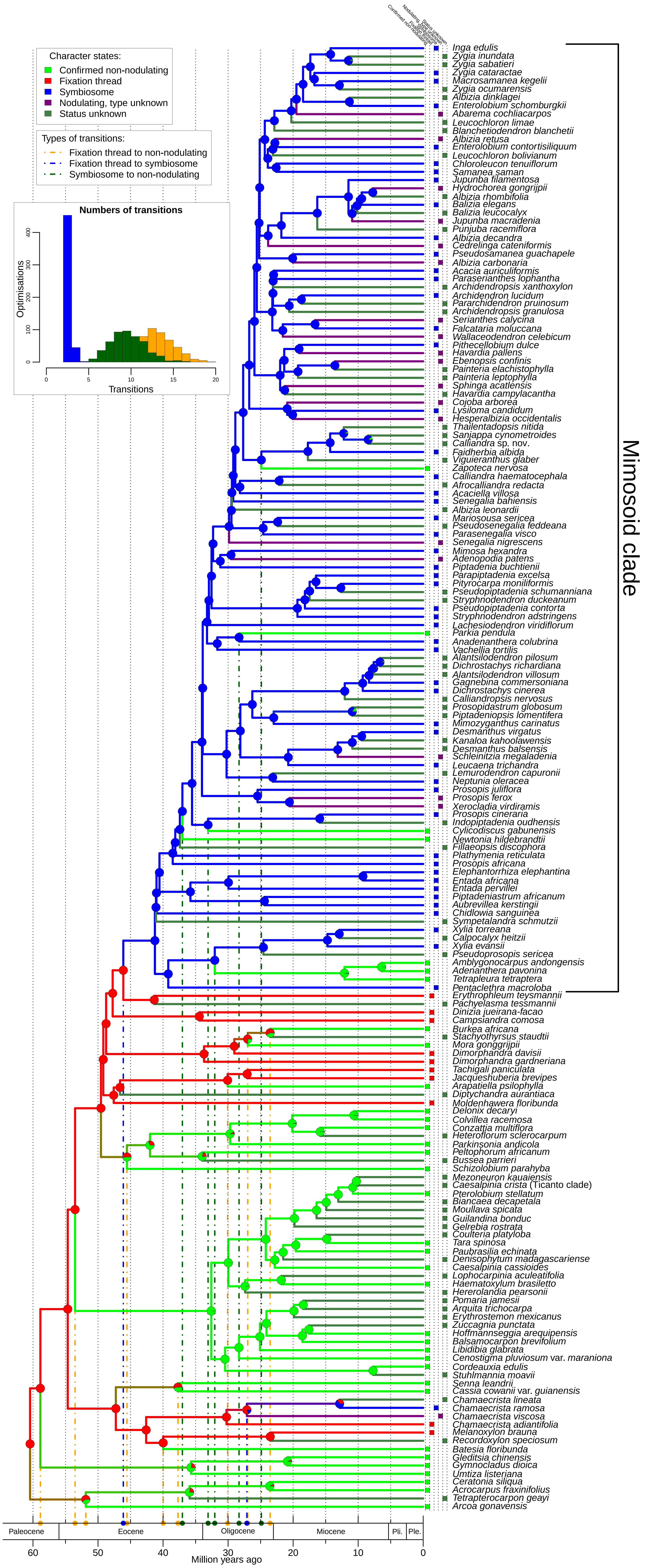
